# Voltage-gated T-type calcium channel blockers reduce apoptotic body mediated SARS-CoV-2 cell-to-cell spread and subsequent cytokine storm

**DOI:** 10.1101/2023.11.03.565419

**Authors:** Thanh Kha Phan, Dylan Sheerin, Bo Shi, Merle Dayton, Liana Mackewicz, Dilara C. Ozkocak, Georgia Atkin-Smith, Nashied Peton, Omar Audi, Rochelle Tixeira, George Ashdown, Kathryn C. Davidson, Marcel Doerflinger, Anna K. Coussens, Ivan K. H. Poon

## Abstract

SARS-CoV-2 typically utilises host angiotensin-converting enzyme 2 (ACE2) as a cellular surface receptor and host serine protease TMPRSS2 for the proteolytic activation of viral spike protein enabling viral entry. Although macrophages express low levels of ACE2, they are often found positive for SARS-CoV-2 in autopsied lungs from COVID-19 patients. As viral-induced macrophage inflammation and overwhelming cytokine release are key immunopathological events that drives exacerbated tissue damage in severe COVID-19 patients, insights into the entry of SARS-CoV-2 into macrophages are therefore critical to understand COVID-19 pathogenesis and devise novel COVID-19 therapies. Mounting evidence suggest that COVID-19 pathogenesis is associated with apoptosis, a type of programmed cell death that often leads to the release of numerous large extracellular vesicles (EVs) called apoptotic bodies (ApoBDs). Here, we showed that ApoBDs derived from SARS-CoV-2-infected cells carry viral antigens and infectious virions. Human monocyte-derived macrophages readily efferocytosed SARS-CoV-2-induced ApoBDs, resulting in SARS-CoV-2 entry and pro-inflammatory responses. To target this novel ApoBD-mediated viral entry process, we screened for ApoBD formation inhibitors and discovered that T-type voltage-gated calcium channel (T-channel) blockers can inhibit SARS-CoV-2-induced ApoBD formation. Mechanistically, T-channel blockers impaired the extracellular calcium influxes required for ApoBD biogenesis. Importantly, blockade of ApoBD formation by T-channel blockers were able to limit viral dissemination and virus-induced macrophage inflammation *in vitro* and in a pre-clinical mouse model of severe COVID-19. Our discovery of the ApoBD-efferocytosis-mediated viral entry reveals a novel route for SARS-CoV-2 infection and cytokine storm induction, expanding our understanding of COVID-19 pathogenesis and offering new therapeutic avenues for infectious diseases.

---

SARS-CoV-2, the causative agent of COVID-19, typically requires the initial binding of its surface spike protein to the surface receptor angiotensin-converting enzyme 2 (ACE2) on host cells. Following ACE2 engagement, the spike protein folds into a virus-host membrane fusion-promoting state through proteolytic cleavage by trypsin-like proteases such as TMPRSS2 (cell surface entry) or cathepsin L (endosomal entry)^1^. Severe SARS-CoV-2 infection is typified by an exacerbated pro-inflammatory ‘cytokine storm’, also known as hypercytokinemia (*e*.*g*., elevated IL-1β, IL-6, TNF) which develops after peak viral titre^2–5^. Macrophages, rapid cytokine producers upon pathogen infection^6^, are a major driver of this COVID-19-associated hypercytokinemia^7,8^. SARS-CoV-2-infected macrophages and macrophage-associated inflammation have been frequently observed in patients’ lung samples^4,9,10^. Despite these clinical observations, macrophages are not permissive to SARS-CoV-2 invasion and replication *in vitro* due to scarce ACE2/TMPRSS2 expression^11–14^. Notably, lack of viral internalisation and/or a low viral replication in macrophages was observed upon SARS-CoV-2 exposure *in vitro*, resulting in limited pro-inflammatory cytokine release^11^. Thus, it remains largely unresolved how SARS-CoV-2 gains entry into macrophages triggering a cytokine storm *in vivo*^5,11,15^.

Apoptotic cell death underpins many important physiological and pathophysiological processes including infection. Dying cells have an active role in their own clearance through the release of ‘find-me’ signals (*e*.*g*. ATP, UTP) to recruit phagocytes to the sites of cell death and the exposure of ‘eat-me’ signals (*e*.*g*. phosphatidylserine) that engage efferocytic receptors^16^. To further aid apoptotic cell clearance, dying cells often break into ‘bite-sized’ (1–5 μm) membrane-bound fragments called apoptotic bodies (ApoBDs) to facilitate efficient dead cell removal via the process known as efferocytosis by tissue-resident professional phagocytes (*e*.*g*., macrophages) or by neighbouring non-professional phagocytes (*e*.*g*. epithelial cells)^17–19^. Recently, we have shown that ApoBD formation is not stochastic as traditionally thought, but rather a highly coordinated process termed apoptotic cell disassembly with three key morphological steps. Firstly, driven by Rho-associated kinase 1 (ROCK1), apoptotic cells form membrane blebs on the cell surface^18,20,21^. The cell membrane then extends, resulting in the generation of long protrusions that radiate the blebs. These protrusions are known as apoptopodia and their formation is negatively regulated by the membrane channel pannexin 1 (PANX1)^17,22^. ApoBDs are then released via the severing of apoptopodia^23^. Notably, ApoBDs can also mediate intercellular communication and elicit a wide range of potent cellular responses in recipient cells^24,25^. Moreover, certain pathogens such as influenza A virus and hepatitis C virus actively induces apoptosis in infected host cells to propagate viral infection to neighboring cells through ApoBD-associated virions^26,27^. However, the role of ApoBD formation in other apoptosis-associated infection settings, particularly during SARS-CoV-2 infection^28–30^, remain largely understudied.

SARS-CoV-2 is known to trigger apoptotic pathways through its accessory viral proteins (ORFs 3a, 6, 7a, 7b, 8a and 9b) and structural proteins (M and N)^31^. In this study, we report that induction of apoptosis by SARS-CoV-2 infection leads to robust formation of ApoBDs that contains infectious virions. Importantly, these ApoBDs released upon SARS-CoV-2 infection can facilitate viral entry into macrophages via an efferocytosis-dependent mechanism. Importantly, SARS-CoV-2-ApoBD-engulfing macrophages but not SARS-CoV-2 exposed macrophages secreted high levels of pro-inflammatory cytokines. Through a drug library screen, we identified the ability of mall molecule inhibitors of voltage-gated T-type calcium channels (T channels) to impair ApoBD formation through blockade of calcium influxes. Of the greatest clinical significance, we further demonstrated that mibefradil, a previously-FDA approved T channel inhibitor effectively dampened ApoBD-efferocytosis-mediated viral entry and inflammation *in vitro* in human macrophages and *in vivo* in a pre-clinical mouse model of severe COVID-19.

## RESULTS

### SARS-CoV-2 infection promotes the formation of ApoBDs that contain infectious virions

SARS-CoV-2 induced apoptosis has been observed in multiple tissues and in circulating T cells in COVID-19 patients and verified *in vitro* in 2D cell culture and organoids, as well as in *in vivo* mouse models^30–35^. Therefore, we first aimed to determine whether ApoBDs are formed upon SARS-CoV-2 infection. In concordance with previous studies, we confirmed that SARS-CoV-2-infected epithelial cells, Vero E6 (**Fig. 1a**) and Calu-3 (**Extended Data Fig. 1a**), underwent apoptosis as measured by staining for caspase 3/7 activation and surface phosphatidylserine exposure using Annexin A5 (A5) (**Extended data Fig. 1b**). Importantly, flow cytometry analysis^36^ showed a large production of ApoBDs from SARS-CoV-2-infected samples at 48 h post-infection, as compared to uninfected controls, to levels equivalent to that achieved with the pro-apoptotic BH3 mimetic drug ABT-737 (**Fig. 1b**).

**Figure 1.**
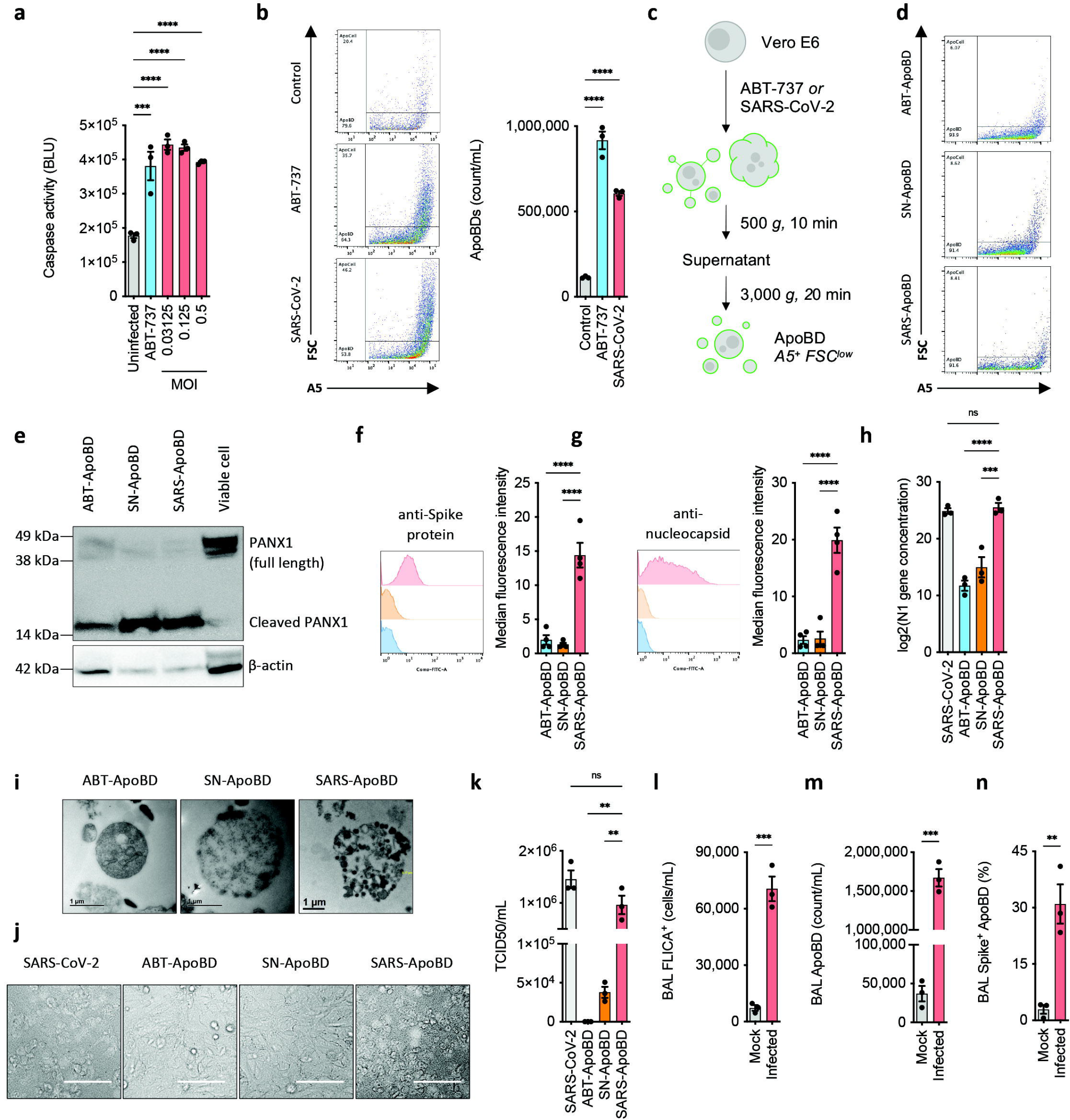
SARS-CoV-2 infection induces formation of infectious virion-containing ApoBDs. **(a)** Detection of apoptotic caspase 3/7 activity in SARS-CoV-2-infected Vero E6 cells by Caspase-Glo®3/7 assay. ABT-737 (1 μM) was included as an apoptosis control. **(b)** Detection of ApoBD formation by flow cytometry according to previous studies^36,37^. **(c)** Schematic for purifying ApoBDs from ABT-737-treated or SARS-CoV-2-infected Vero E6 cells. **(d)** ApoBD purity assessment by flow cytometry analysis; ABT-ApoBD (ApoBDs derived from ABT-737 treatment, i.e., non-infectious ApoBD control), SN-ApoBD (ApoBDs incubated with SARS-CoV-2 supernatant, i.e., a control for residual and/or sticky viruses) and SARS-ApoBD (ApoBDs from SARS-CoV-2 infected cells). **(e)** Detection of caspase-cleaved substrates, PANX1, as ApoBD markers by immunoblotting. (**f)** Viral Spike protein, **(g)** viral nucleocapsid protein, **(h)** viral N1 RNA and **(i)** intact virions were detected using flow cytometry, qPCR and transmission electron microscopy, respectively. Infectivity of ApoBDs were determined by **(j)** bright-field imaging (scale bar, 100 μm) and **(k)** TCID50 assay. **(l)** Apoptotic cells (FLICA staining), **(m)** ApoBD formation (TO-PRO-3 and A5 staining) and **(n)** spike^+^ ApoBDs (with anti-spike protein antibody) were measured in murine bronchioalveolar lavage using flow cytometry. Bronchioalveolar lavage were collected from mice inoculated intranasally with mouse-adapted SARS-CoV-2 strain at 3 d.p.i. Data are mean±S.E.M of n≥3. Data in (i) and (j) are representative of 3 independent experiments. ^*^ denotes p-value of <0.05; ^**^ p<0.01, ^***^ p<0.001, ^****^ p<0.0001, ns: not significant, as determined by Student’s t-test or one-way ANOVA with a Tukey post-hoc test.

Since ApoBDs and other types of EVs reportedly can harbour and traffic viral materials and/or intact virion, we isolated ApoBDs derived from SARS-CoV-2-infected Vero E6 supernatant (denoted ‘SARS-ApoBD’) using our previously established differential centrifugation approach^37^ (**Fig. 1c**) to determine its biomolecular contents. Of note, the differential centrifugation approach consistently enabled ApoBD isolation to 90-95% purity (**Fig. 1d**). For comparison, we also isolated non-infectious control ApoBDs generated from ABT-737 treatment (denoted ‘ABT-ApoBD’) and to control for any residual SARS-CoV-2 virions present in the media of the SARS-ApoBD preparation, we also incubated ABT-ApoBD with the leftover supernatant following SARS-ApoBD isolation (denoted ‘SN-ApoBD’). The ApoBD identity of each of these preparations were further verified by the presence of ApoBD-specific protein markers^38^ such as caspase-cleaved PANX1 (**Fig. 1e**). In contrast to both ApoBD controls, we confirmed SARS-ApoBDs carried SARS-CoV-2 spike protein (**Fig. 1f**), nucleocapsid protein (**Fig. 1g**) and viral nucleocapsid RNA (**Fig. 1h**). Importantly, using transmission electron microscopy whole virions were detected abundantly within SARS-ApoBDs, minimally on the surface of SN-ApoBDs and absent from ABT-ApoBDs (**Fig. 1i**). The presence of viral materials and whole virions in ApoBDs generated from SARS-CoV-2 infected cells suggests their ability to traffic infectious virions. Through bright-field imaging (**Fig. 1j**) and viral load quantification by 50% tissue culture infectious dose (TCID50) assay (**Fig. 1k**), comparable cytopathic effect was observed for Vero E6 cells incubated with SARS-ApoBDs and those infected directly with SARS-CoV-2, but to a much lesser extent with SN-ApoBD and ABT-ApoBD controls. Together, these data confirm ApoBDs generated from SARS-infected cells can harbour infectious virions.

To examine whether virus-carrying ApoBDs are generated *in vivo*, we infected aged mice with a mouse-adapted SARS-CoV-2 strain that could elicit a severe form of COVID-19^39^. In bronchoalveolar lavage (BAL) samples collected at day 3 post-infection by flow cytometry (**Extended data Fig. 1c**), apoptotic cells were shown to be significantly elevated compared to mock infected controls, as indicated by an elevated number of cells with active caspase 3/7 (**Fig. 1l**). Concomitantly, a significantly elevated ApoBD level were also detected in the infected mouse BAL, as compared to mock control (**Fig. 1m**). Strikingly, 30-40% of those ApoBDs contain viral spike protein (**Fig. 1n**).

### SARS-CoV-2 infection-derived ApoBDs facilitate viral entry and inflammation in macrophages via efferocytosis-mediated viral entry

Given that macrophages are key drivers of COVID-19 hypercytokinemia and are identified *in vivo* to contain virus^5,11,15^ despite low ACE2 expression^7,8^ (**Extended data Fig. 2a**), we sought to examine the potential role of virus-carrying ApoBDs in facilitating viral entry and hyperinflammatory processes in macrophages. Firstly, co-culturing human monocyte-derived macrophages (MDMs) with SARS-ApoBD for 48 h, we detected viral spike proteins in macrophages, by fluorescence microscopy and flow cytometry analysis but not when co-cultured with either control ApoBDs (ABT-ApoBDs or SN-ApoBDs) or SARS-CoV-2 alone (**Fig. 2a,b**). qPCR analysis also revealed significant detection of N1 RNA in SARS-ApoBD-co-cultured macrophages (**Fig. 2c**). Furthermore, SARS-ApoBD-co-cultured macrophages was the only condition that resulted in a significant decrease in macrophage viability, further indicating viral entry (**Fig. 2d,e**). Of note, performing a TCID50 assay using the resulting macrophage-derived conditioned media revealed limited infectious virions produced by the macrophages, indictive of aborted infection (**Extended data Fig. 2b**). Critically, SARS-ApoBD-treated MDMs released significant levels of severe COVID-19 associated cytokines TNF, IL-6, IL-1β, IL-10 and IL-8 (**Fig. 2f**). These effects were not observed with direct SARS-CoV-2 or control ApoBDs co-culture (**Fig. 2f**), further suggesting a specific attribution to virus-carrying ApoBDs engulfed by macrophages, rather than free/co-pelleted viruses or ApoBDs alone.

**Figure 2.**
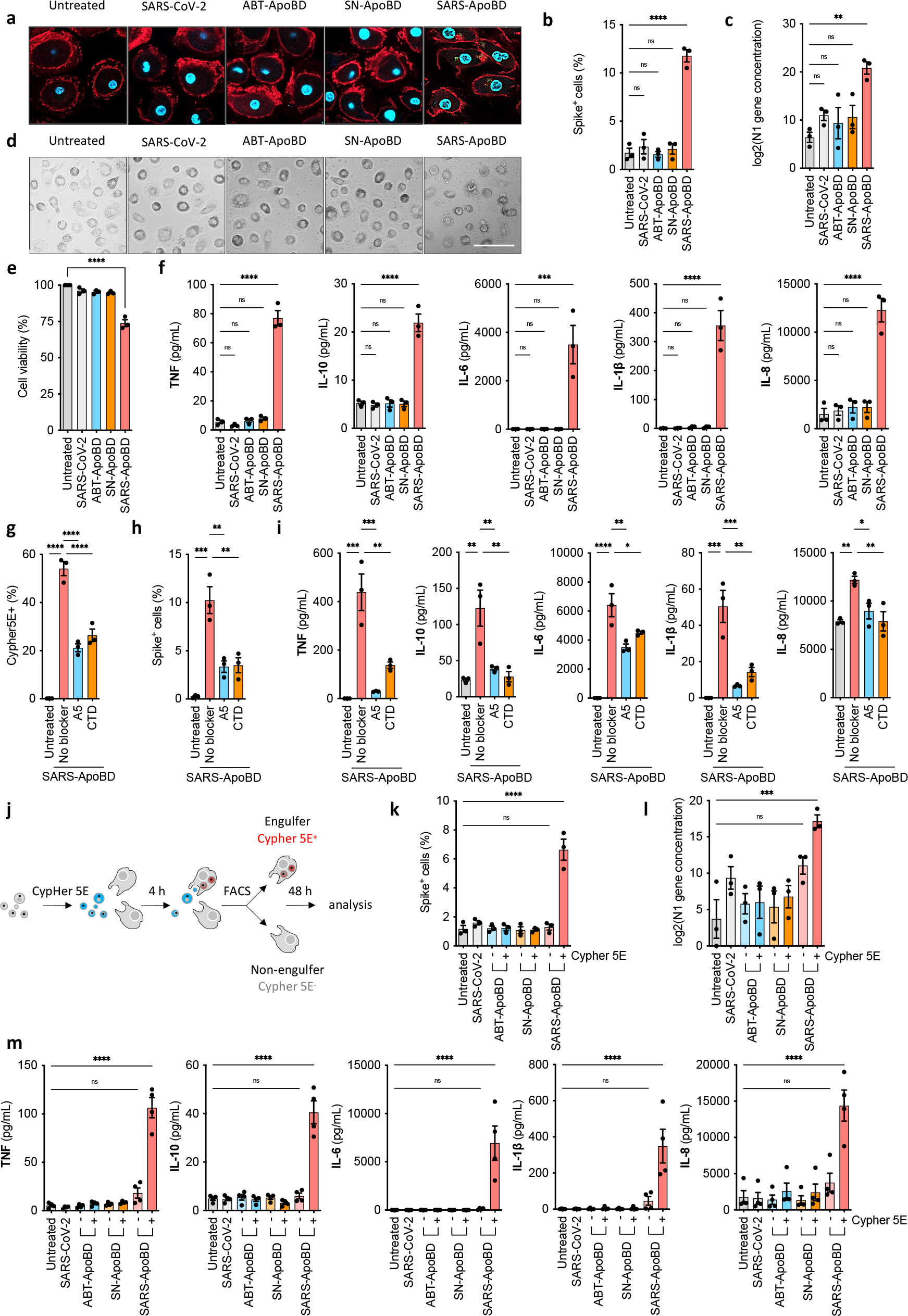
SARS-ApoBDs induce inflammation in macrophages via efferocytosis-mediated viral uptake. **(a)** Immunofluorescence imaging showing nucleus staining (light blue), F-actin (red) and viral Spike protein (green). **(b)** Viral spike-staining flow cytometry and **c)** N1 RNA-targeting qPCR analyses for ApoBD-co-incubated human MDMs (MDM:ApoBD ratio of 1:3). Cytopathic effect on MDMs at 48 post-co-incubation were determined by **(d)** bright-field imaging (scale bar, 100 μm) and **(e)** cell viability assay. **(f)** Inflammatory cytokines released by ApoBD-incubated MDM, detected using bead-based cytokine assay. **(g)** Efferocytosis of SARS-CoV-2 infection-derived ApoBDs by MDMs and inhibition thereof by A5 and cytochalasin D, indicated by CypHer5E staining. **(h)** Reduced spike protein detection and **(i)** impaired cytokine release in SARS-ApoBD-treated MDM by efferocytosis inhibitors. **(j)** Schematic for the FACS of ApoBD engulfing MDMs (CypHer5E^+^) and non-engulfing MDMs (CypHer5E^-^), followed by detection of viral uptake, as determined by **(k)** spike protein signal and **(l)** N1 RNA, as well as **(m)** cytokine release. Data are mean±S.E.M of n≥3. Data in (a) and (d) are representative of 3 independent experiments. ^*^ denotes p-value of <0.05; ^**^ p<0.01, ^***^ p<0.001, ^****^ p<0.0001, ns: not significant, as determined by Student’s t-test or one-way ANOVA with a Tukey post-hoc test.

We next sought to confirm ApoBD-mediated SARS-CoV-2 entry into macrophages was mediated through efferocytosis of ApoBDs. An engulfment assay using the pH-dependent CypHer5E stain as an engulfment indicator showed that MDMs readily efferocytosed SARS-ApoBDs (**Fig. 2g**). By contrast, treatment with A5 to sequester the ‘eat-me’ signal phosphatidylserine on ApoBDs or cytochalasin D (CTD), an inhibitor of actin polymerisation required for efferocytosis, significantly impaired SARS-ApoBD engulfment (**Fig. 2g**). Blockade of SARS-ApoBD uptake by A5 or CTD treatment also reduced the proportion of spike^+^ MDMs (**Fig. 2h**) and markedly reduced levels of macrophage cytokine release (**Fig. 2i**). Next, we isolated SARS-ApoBD engulfing and non-engulfing MDMs by fluorescence activated cell sorting (FACS) based on high and low CypHer5E signal, respectively (**Fig. 2j**), with SARS-ApoBD-engulfing MDMs (CypHer5E^+^) showed signs of viral uptake as evidenced by an increase in the percentage of spike^+^ cells (**Fig. 2k**) and high level of viral N1 RNA (**Fig. 2l**), as well as substantial cytokine production (**Fig. 2m**). In contrast, their non-engulfing (CypHer5E^-^) counterparts and MDMs that engulfed ABT-ApoBD or SN-ApoBD did not show these effects (**Fig. 2k-m**). These data collectively demonstrate that efferocytosis of ApoBDs is necessary for ApoBD-mediated SARS-CoV-2 entry and subsequent triggering of macrophage hypercytokinemia.

### Mibefradil and other voltage-gated T-type calcium channel blockers impair SARS-CoV-2-induced ApoBD formation

Disassembly of apoptotic cells into ApoBDs promotes clearance of apoptotic materials and genetic disruption of ApoBD biogenesis leads to noticeable reduction in efferocytosis efficiency^18^. Thus, we reasoned that identification of small molecules that could reduce infection-induced ApoBD formation could subsequently limit ApoBD-efferocytosis-mediated SARS-CoV-2 entry into macrophages and associated inflammation. To this end, we tested inhibitors of ApoBD formation based on our previous drug screening on apoptotic Jurkat T cells^40^, a well-established model for apoptotic cell disassembly^18,36,41^, prior to testing on an infection model^18^ (**Fig. 3a**). In this model, trovafloxacin (TVF) was added to inhibit the key negative regulator of apoptotic cell disassembly, PANX1, to maximise the generation of ApoBD formation by Jurkat T cells^18,36,41^, hence allowing the identification for potent ApoBD formation inhibitors. From this screen, a number of calcium channel blockers in the drug library showed certain inhibitory effects on apoptotic cell disassembly (**Extended data Fig. 3a**). In a secondary, validating screen, we found the calcium channel blocker mibefradil (MBF) substantially impaired ApoBD formation in a dose-dependent manner (**Fig. 3b**). As shown by time-lapse differential interference contrast (DIC) microscopy, MBF treatment attenuated the protrusion and fragmentation steps of ApoBD biogenesis, but not membrane blebbing, during the progression of apoptosis (**Fig. 3c and Supplementary videos 1a−b**).

**Figure 3.**
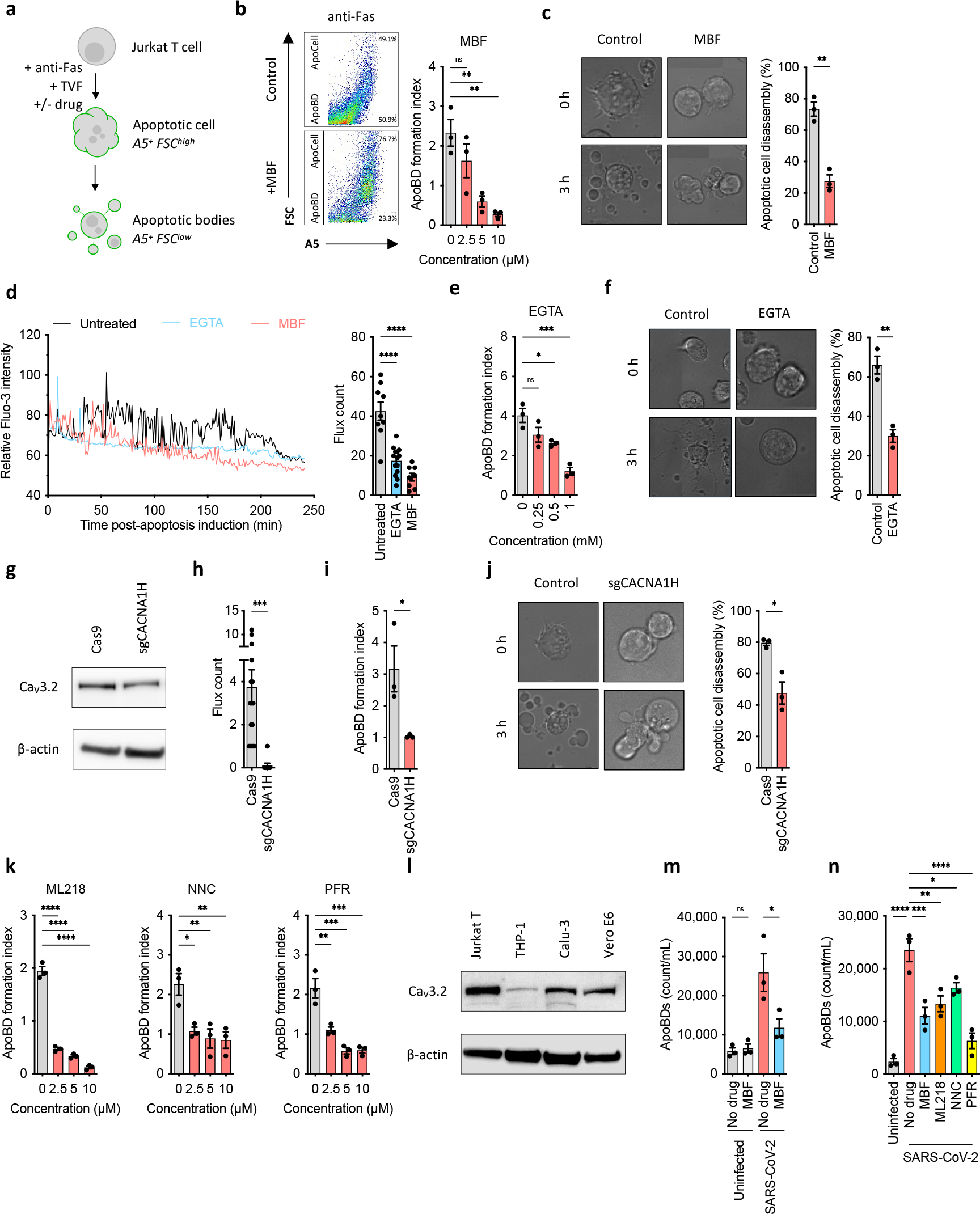
Voltage-gated T-type calcium channel blockers, novel inhibitors of apoptotic cell disassembly, impair SARS-CoV-2-induced ApoBD formation. **(a)** Schematic for the LOPAC® 1280 drug library screening to identify novel inhibitors of apoptotic cell disassembly. Trovafloxacin, a known apoptotic cell disassembly enhancer, was added to maximise ApoBD formation, hence allowing the screening for potent, robust blockers **(b)** Detection of ApoBD formation following MBF treatment by flow cytometry using TO-PRO-3 and A5 staining. **(c)** Time-lapse DIC microscopy of anti-Fas (1:1000)-treated Jurkat T cells with or without 10 μM mibefradil (MBF). The level of apoptotic cell disassembly was quantified as the percentage of cells undergoing fragmentation **(d)** Time-lapse Fluo-4 signal intensity (left panel) and peak counts (right panel), indicative of calcium influxes, of 1 mM EGTA-, 10 μM mibefradil- or vehicle-treated THP-1 monocytes post-UV irradiation, as measured by confocal imaging. **(e)** Detection of ApoBD formation following EGTA treatment by flow cytometry. **(f)** Time-lapse DIC microscopy to apoptotic cell disassembly in the presence of 1 mM EGTA or vehicle control. **(g)** Immunoblotting showing the knockdown of CACNA1H gene encoding for Ca_V_3.2 in Jurkat T cells using CRISPR/Cas9 gene editing. **(h)** Calcium influxes, indicted by Fluo-4 peak counts, of CACNA1H-deficient cells and Cas9 control. **(l)** Flow cytometry analysis and **(j)** Time-lapse DIC microscopy to quantify and monitor ApoBD formation in CACNA1H-knockdown cells or Cas9 control following apoptotic induction using anti-Fas. **(k)** Flow cytometry analysis of anti-Fas-treated Jurkat T cells treated with other T-channel inhibitors, including ML218, NNC 55-0396 and penfluridol. **(l)** Immunoblotting of CaV3.2 in various cell lines including epithelial Calu-3 and Vero E6, the commonly used host cell models for SARS-CoV-2 infection. Detection of SARS-CoV-2 infection-induced ApoBD formation following treatment with **(m)** mibefradil and **(n)** other T-channel inhibitors. Data are mean±S.E.M of n≥3. Data in (c), (f), (g), (h), (j) and (i) are representative of 3 independent experiments. ^*^ denotes p-value of <0.05; ^**^ p<0.01, ^***^ p<0.001, ^****^ p<0.0001, ns: not significant, as determined by Student’s t-test or one-way ANOVA with a Tukey post-hoc test.

MBF is a T-type voltage-gated calcium channel (T-channel) blocker that inhibits T-channel-mediated calcium influx from the extracellular space into the cytosol^42^, previously FDA-approved for the treatment of hypertension and angina. To better understand the mechanism underpinning the inhibition of ApoBD formation by MBF, we performed calcium imaging by confocal microscopy using the calcium probe Fluo-4. MBF was found to impair both the intensity and the number of calcium influxes detected post-apoptosis induction and prior to apoptotic cell fragmentation (**Fig. 3d** and **Supplementary videos 2a−c)**. Similar inhibitory effect was also observed upon extracellular calcium chelation using EGTA (**Fig. 3d**). Remarkably, EGTA treatment also inhibited ApoBD formation (**Fig. 3e**), and the formation of membrane protrusion and cell fragmentation during progression of apoptosis (**Fig. 3F**). It is worth noting that the observed reduction in ApoBD formation following MBF and EGTA treatment was not due to impaired apoptosis induction as neither MBF nor EGTA altered the levels of cell death induced by anti-Fas (**Extended Data Fig. 3b**). These observations suggest MBF acts on calcium influxes that drive ApoBD biogenesis.

Among the three T-channel isoforms (CaV3.1, CaV3.2 and CaV3.3 encoded by *CACNA1G, CACNA1H* and *CACNA1I* respectively), our model Jurkat T cells express predominantly CaV3.2 (**Extended Data Fig. 3**). Next, to deduce the relevance of T-channels in apoptotic cell disassembly, we attempted to generate a CaV3.2-deficient cell line using CRISPR/Cas9. Although we were unable to generate a stable knockout cell line, the resultant CaV3.2 knockdown cells (**Fig. 3h**) exhibited impaired calcium influxes post-apoptosis induction (**Fig. 3i** and **Supplementary videos 3a−b**) as well as decreased ApoBD formation due to impaired protrusion formation and fragmentation (**Fig. 3j−k** and **Supplementary videos 4a−b**). In addition, other T-channel specific inhibitors including NNC 55-0396 (NNC) (identified in the initial screen, **Extended Data Fig. 3a**), ML218, and penfluridol (PFR) exerted similar dose-dependant inhibitory effects on ApoBD formation (**Fig. 3l**). By contrast, blockers of other types of calcium channels including L-type calcium channel (nifedipine and nicardipine), N-type calcium channel (PD173212), calcium-release-activated calcium channel protein 1 Orai1 (AnCoA4) or calcium storage release ryanodine receptor (dantrolene) or inositol trisphosphate receptor (2-aminoethoxydiphenylborane) did not inhibit ApoBD formation (**Extended Data Fig. 3d**). Therefore, these data indicate that T-channels, the targets of MBF, are positive regulators of apoptotic cell disassembly. Of note, T-channel isoforms such as CaV3.2 are expressed in many tissues and cell lines, including epithelial Calu-3 and Vero E6 cells (**Fig. 3m**), suggesting their druggability to block SARS-CoV-2 infection-induced ApoBD formation. Comparable to the Jurkat T cell model, MBF treatment markedly inhibited the formation of ApoBDs by SARS-CoV-2 infected Vero E6 cells (**Fig. 3n**) without affecting the levels of apoptosis or viral infectivity (**Extended Data Fig. 3e-f**). Similarly, the other T-channel inhibitors NNC, ML218 and PFR also reduced ApoBDs release from SARS-CoV-2 infected apoptotic cells (**Fig. 3o**).

### Mibefradil reduces SARS-CoV-2 uptake and inflammation in macrophages *in vitro* and *in vivo*

Inhibition of SARS-CoV-2 infection-derived ApoBD formation by T-channel inhibitors such as MBF entails their potential to limit ApoBD-efferocytosis-mediated SARS-CoV-2 entry into macrophages and dampen subsequent inflammation. To this end, we treated SARS-CoV-2 infected Vero E6 cells with or without MBF for 48 h and then transferred the free virus-depleted conditioned media (CM) onto MDMs (**Fig. 4a**). CM from MBF-treated cells resulted in significantly reduced the levels of ApoBD efferocytosis measured by Cypher5E+ and viral entry as indicated by viral N1 RNA in MDMs compared to vehicle control CM (**Fig. 4b,c**). Significantly, MDMs cultured in MBF-treated CM also showed enhanced viability (**Fig. 4d-e**) and reduced cytokine secretion (**Fig. 4f**). Finally, we tested the effect of MBF on SARS-CoV-2 infected aged mice that we previously demonstrated develop a severe form of COVID-19^39^. MBF was intranasally administered on day 1 and day 2 post infection, with viral peak occurring on day 2 in this model and cytokine peak on day 3 (**Fig. 4g**). Analogous to the *in vitro* findings (**Fig. 3n**), the number of ApoBDs detected in BAL samples of MBF-treated mice on day 3 post-infection was significantly lower than that of vehicle control (**Fig. 4h, Extended Data Fig. 4**). This effect also aligned with a reduction in spike^+^ alveolar macrophages in the BAL, suggesting less viral entry into macrophages *in vivo* upon MBF administration (**Fig. 4i**), as well as significantly reduced viral load as shown by the lung TCID50 assay (**Fig. 4j**) and reduced TNF and IL-6 secretion (**Fig. 4k)**. In terms of innate immune cell infiltration into the lung, MBF treatment dramatically reduced the levels of infiltrating macrophages, neutrophils and, monocytes (**Fig. 4l**, with near amelioration of the pulmonary pathology indicative of severe COVID-19 upon histological examination (**Fig. 4m**).

**Figure 4.**
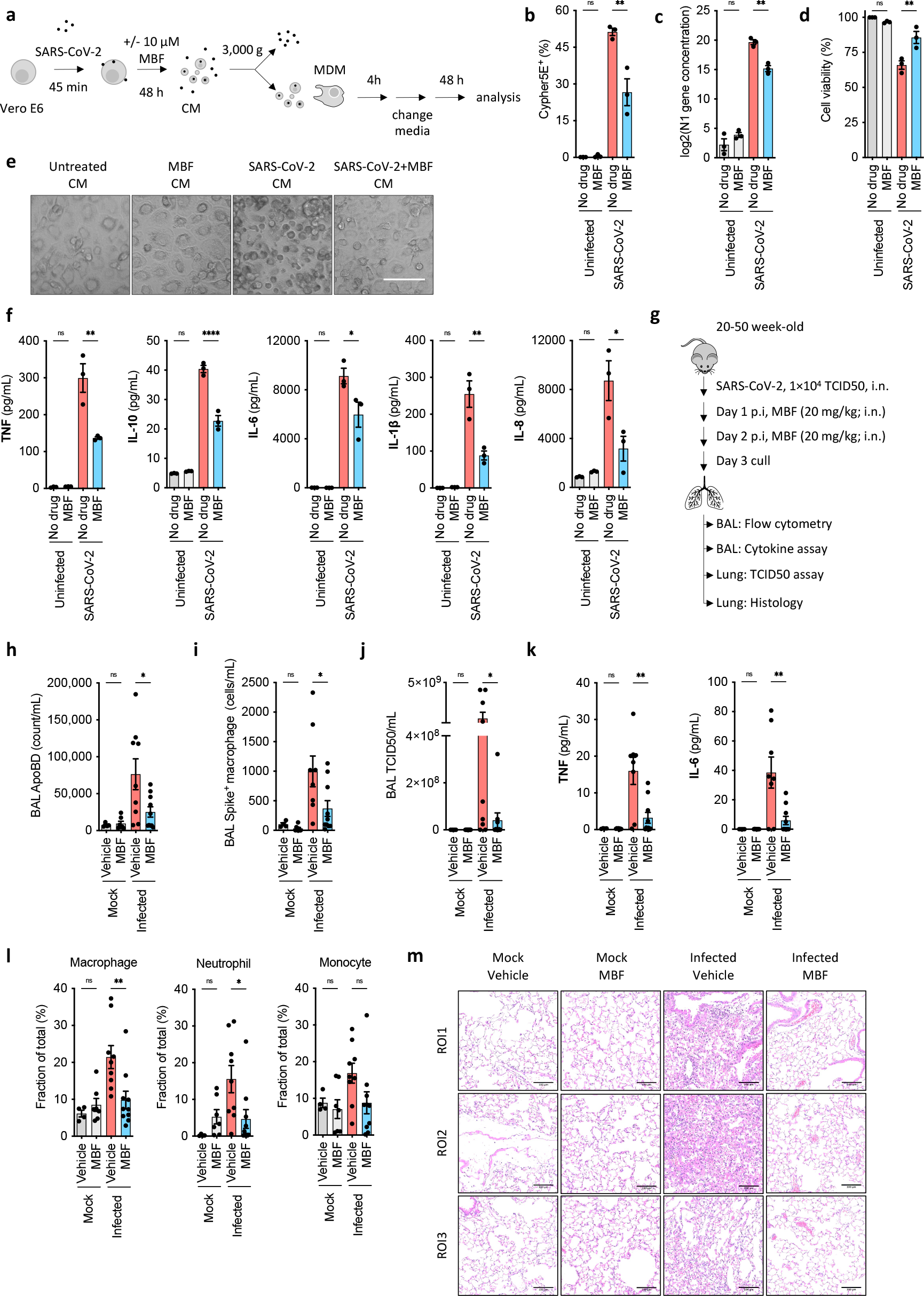
Mibefradil reduces SARS-CoV-2 entry and inflammation *in vitro* and *in vivo*. **(a)** Schematic for transfer of condition media (CM) to elucidate the effect of MBF on viral uptake, cytopathic effect and inflammatory responses by MDMs. **(b)** Flow cytometry analysis of MDMs efferocytosing apoptotic materials in condition media, as measured by CypHer5E positivity. **(c)** qPCR analysis, **(d)** cell viability assay as well as **(e)** bright-field imaging and **(f)** cytokine bead array to determine viral N1 RNA, cytopathic effect and inflammation in MDMs upon condition media treatment. **(g)** Schematic for mibefradil administration in SARS-CoV-2-infected mice and post-treatment analysis to detect **(h)** ApoBD formation (by flow cytometry of the bronchioalveolar lavage), **(i)** spike^+^ macrophages (BAL), **(j)** viral load (by TCID50 assay on lung samples), **(k)** inflammatory cytokines (by cytokine bead array of BAL), **(l)** immune cell infiltration (BAL), and **(m)** lung histology (H&E staining). Data in b, (c), (d) and (f) are mean±S.E.M of n≥3. Data in (e) are representative of 3 independent experiments. Data in (h−l) are mean±S.E.M of n≥5. Data in (m) are representative of at least 5 mice per group. ^*^ denotes p-value of <0.05; ^**^ p<0.01, ^***^ p<0.001, ^****^ p<0.0001, ns: not significant, as determined by Student’s t-test or one-way ANOVA with a Tukey post-hoc test.

## DISCUSSION

The intricate mechanisms by which SARS-CoV-2 enters and interacts with host cells have been the subject of extensive research since the start of the COVID-19 pandemic, contributing to the growing understanding of viral pathogenesis. Nevertheless, how SARS-CoV-2 enters macrophages and drives detrimental hypercytokinemia in COVID-19 patients is yet to be fully defined. Macrophages are heterogeneous and the expression of key host factors for SARS-CoV-2 entry, *ACE2* and *TMPRSS2*, are low to undetectable in human MDMs and BAL macrophages investigated herein. Consistent with this notion, recent *in vitro* studies showed that viral internalisation and/or viral replication in human MDM upon SARS-CoV-2 exposure is limited even at high MOI of 20, and was unable to trigger inflammation^43^. However, it should be noted that in the presence of anti-viral spike antibodies, ACE2-negative macrophages can take up antibody-opsonised SARS-CoV-2 through FcγR receptors and causes systematic inflammatory responses despite non-productive abortive infection^44^. Here, we describe that in the absence of opsonizing antibodies and at a relative low MOI of 3, SARS-CoV-2 can also enter into ACE2-negative/low human primary macrophages through the efferocytosis of ApoBDs released by apoptotic SARS-CoV-2 infected epithelial cells. Unlike SARS-CoV-2 exposure alone, human MDM became infected and was able to elicit secretion of TNF, IL-6, IL-1β, IL-8 and IL-10 indicative of the cytokine storm in severe COVID-19 patients^45,46^. Therefore, our findings suggest a potential alternative mechanism for viral entry and exacerbated macrophage-driven inflammation in COVID-10 patients. Although we deduced that efferocytic machineries underpin this viral entry pathway, exact efferocytic receptors and details of downstream viral escape pathway warrants further research. As ApoBDs are readily circulating and efficiently engulfed by non-professional phagocytes such as endothelial and epithelial cells, our findings herein have potential to be extended into other pathophysiological hallmarks of SARS-CoV-2 infection including, but not limited to, multi-organ injuries, cardiovascular damages, and neurological complications.

Small EVs can exert a variety of effects on recipient cells, often mediated through their accompanying cargoes^47^. Notably, circulating EVs of 50−500 nm (classified as small EVs^48^) in COVID-19 patients can be endocytosed and cause apoptosis in recipient pulmonary endothelial cells^49^. However, whether such EVs can facilitate viral entry and impact inflammation has not been determined. Here, in line with existing literature, our study confirmed SARS-CoV-2-induced apoptosis in various host cells as well as in the murine infection model, resulting in the formation of SARS-CoV-2-carrying ApoBDs (classified as large EVs) that are released exclusively during apoptosis. Importantly, we demonstrated that the engulfment of SARS-CoV-2-carrying ApoBDs can subsequently aid viral entry into macrophages and promotes inflammation despite abortive infection. Our observation raises several opportunities to target this pathway of viral entry into macrophages. This include (i) targeting the induction of apoptosis, (ii) the formation of ApoBDs, and (iii) the efferocytosis of ApoBDs. In fact, blockade of apoptosis through caspase-6 inhibition arrests viral replication and substantially attenuates lung pathology in golden Syrian hamsters infected with SARS-CoV-2-infected golden^50^. Alternatively, targeting the efferocytosis machinery may also be another approach to block viral entry into macrophages and prevent hyperinflammation. However, it is worth noting that cell clearance itself is also an important mechanism to limit inflammation^51^. As the molecular basis of ApoBD formation is yet to be fully elucidated it thus represents novel opportunities for therapeutic targeting. We identified T-channels as new regulators of apoptotic cell disassembly and demonstrated the blockade of T-channels with MBF can limit SARS-CoV-2-induced ApoBD formation through the blockade of calcium influxes. These inhibitors were shown to mitigate ApoBD-mediated viral entry and not only dampen hypercytokinemia in primary human macrophages and mouse BAL macrophages but also ameliorate overall lung inflammation. As only a few enzyme-inhibiting antivirals have been approved with greatest efficacy prior to viral peak and limitedly prescribed for COVID-19, new/repurposed drugs with different mode of mechanisms like MBF (highly tolerable, topically applicable, efficacy when given during viral peak) not only reduce disease burden in severe patients, but also lower the risk of drug resistance and have a longer efficacy window. Together, our findings not only provide further insights into SARS-CoV-2 pathogenesis but rationalise the development of ApoBD formation-targeting drugs to tackle severe COVID-19 as well as other apoptosis-inducing newly emerging viral threats.

## METHODS

### Mammalian cell cultures

#### Cell lines

Monkey kidney epithelial cells Vero E6 (clone CCL81) and human lung epithelial cells Calu-3 were respectively cultured in DMEM medium and EMEM medium (ThermoFisher) supplemented with 10% heat-inactivated fetal bovine serum (FBS). Human Jurkat T cells and human THP-1 monocytes were cultured in RPMI 1640 medium (ThermoFisher), supplemented with 10% FBS, penicillin (50 U/ml), streptomycin (50 μg/ml) and MycoZap (Lonza, Switzerland). All cells are cultured at 37 °C in a humidified atmosphere with 5% CO_2_.

#### Human monocyte-derived macrophage preparation

Human monocyte-derived macrophages (MDMs) were prepared from fresh buffy coat obtained from the Australian Red Cross Lifeblood Service with donor consent as previously described^52,53^. Appropriate ethics approval was granted by the La Trobe University and the Walter and Eliza Hall Institute of Medical Research (WEHI) Human Ethics Committees (WEHI HREC #18_09LR). Briefly, peripheral blood mononuclear cells were isolated from buffy coats by density gradient centrifugation with Lymphoprep (StemCell Technologies). CD14^+^ monocytes were then obtained by positive magnetic isolation using human CD14 microbeads (Miltenyi Biotec) and a MACS LS column with a QuadroMacs™ separator (Miltenyi Biotech). Monocytes were differentiated into GM-CSF-polarised (M1) MDMs over seven days in RPMI 1640 media supplemented with 1 mM L-glutamine (Gibco), 2 mM sodium pyruvate, 10% human adult AB serum (Sigma-Aldrich) and 5 ng/mL GM-CSF (Miltenyi Biotec) at 37 °C in a humidified atmosphere with 5% CO2.

#### Human neutrophil preparation

Human neutrophils were prepared from fresh Sodium Heparin blood obtained from consenting donors through the WEHI volunteer blood donor registry as previously described^54^. Ethics approval was granted by WEHI HREC #18_09LR. Whole blood was diluted 1:1 with 1× PBS (Thermo Fisher) and layered over Lymphoprep followed by density gradient centrifugation. Neutrophils were isolated from the cell pellet following red blood cell (RBC) lysis (8% Ammonium chloride [NH4Cl], 0.8% Sodium bicarbonate [NaHCO3] and 0.4% Ethylenediaminetetraacetic acid [EDTA]). Following two washes in PBS, neutrophils were resuspended in OptiMEM (Thermo Fisher) and 1 x 10^7^ cells were seeded evenly into 3 wells of a 6-well cell culture plate (Nest Scientific). Cells were incubated for 6h at 37°C in a humidified atmosphere with 5% CO2. SN was removed and 1 ml Trizol added to each well, cells homogenised and Trizol stored at -80°C before RNA extraction.

### Mice

Aged (20-50 weeks old), female wild-type C57BL/6J mice were bred and maintained in the Specific Pathogen Free Physical Containment Level 2 Bioresources Facility at WEHI. All procedures involving animals and live SARS-CoV-2 strain were conducted in an OGTR-approved Physical Containment Level 3 (PC3) facility at WEHI (Cert-3621). Mice were transferred to the PC3 laboratory for all SARS-CoV-2 infection experiments at least 4 days prior to experiments. All mouse strains and procedures were reviewed and approved by the WEHI Animal Ethics Committee and were conducted in accordance with the Prevention of Cruelty to Animals Act (1986) and the Australian National Health and Medical Research Council Code of Practice for the Care and Use of Animals for Scientific Purposes (1997).

### SARS-CoV-2 strains, viral infection and mibefradil treatment

#### Infection of mammalian cells

Mammalian cells were infected with a clinical isolate of the originating Wuhan strain of SARS-CoV-2 (hCoV-19/Australia/VIC01/2020, obtained from the Victorian Infectious Disease Reference Laboratory). To prepare viral stocks, viruses were grown in Vero E6 cells in serum-free DMEM media supplemented with 1 μg/mL TPCK-treated trypsin (Thermo Fisher). Conditioned media were collected after 48 h and centrifuged at 3,000 g for 10 minutes before passing through a 0.22 μm Millex® syringe filters (Merck Millipore). For viral infection, virus stocks were diluted in serum-free media (with 1 μg/mL TPCK-treated trypsin) to the indicated multiplicity of infection (MOI) and added onto mammalian cells. After 1 h incubation, infection media were removed and replaced with respective culture media supplemented with 2% FBS and further incubated for 48 h.

#### Infection of mouse models

For SARS-CoV-2 murine infection, the mouse-adaptive SARS-CoV-2 VIC2089 P21 isolate (generated through serial passaging of the clinical isolate hCoV-19/Australia/VIC2089/2020)^39^ was used. Briefly, virus stocks were diluted in serum-free DMEM to a final concentration of 1×10^4^ TCID50/mouse and inoculated intranasally (30 μL) into methoxyflurane-anesthetised mice. After infection, animals were visually checked and weighed daily before euthanised at 3 d.p.i.

For intranasal administration of mibefradil, female mice (20-50 weeks old, age-matched) were infected with SARS-CoV-2 (1×10^4^ TCID50/mouse) or mock. MBF (reconstituted in 1× PBS, 20 mg/kg) or PBS vehicle were administered intranasally on 1 and 2 d.p.i. After infection, animals were visually checked and weighed daily before euthanised at 3 d.p.i.

### ApoBD detection

#### ApoBD detection in conditioned media

ABT-737-treated or SARS-CoV-2-infected cells (detached using Trypsin/EGTA) and their respective conditioned media were combined and stained with fluorescently-labelled annexin A5 (BD Biosciences, 1:2,000) and 0.2 μM TO-PRO-3 (Life Technologies) in 1× A5 binding buffer (BD Biosciences) at room temperature for 10 minutes. Samples are immediately placed on ice and SPHERO™ AccuCount beads (Spherotech) were added prior to analysis using a BD FACSAria™ Fusion flow cytometer and FACSDiva 6.1.1 software (BD Bioscience). Data were analysed using FlowJo 10.7.1 software (Tree Star), as previously described^55^.

#### ApoBD detection in BAL samples

Mock or SARS-CoV-2-infected mice were humanely euthanised at 3 d.p.i. To collect bronchioalveolar lavage (BAL), the trachea was cannulated and 4× 0.5 mL aliquots of 1× PBS was delivered to and captured from the lungs. BAL samples were then centrifuged at 3,000 *g* for 20 min to collect cells and ApoBDs for staining with a combination of SR-DEVD-FMK (caspase 3/7 activity detection dye, Thermo Fisher), annexin A5-V450 and TO-PRO-3 in 1× A5 binding buffer at room temperature for 10 min. Following fixation (with 2% paraformaldehyde) and permeabilisation (using 0.5% saponin), cell/ApoBD samples were incubated with FITC-conjugated anti-viral spike antibodies (1:250, RnD Systems). SPHERO™ AccuCount beads (Spherotech) were also added prior to flow cytometry analysis using a BD FACSAria™ Fusion flow cytometer.

#### ApoBD detection in drug treatments

To induce apoptosis and ApoBD formation in Jurkat T cells, cells at 1.5×10^6^ cells/mL in serum-free RMPI 1640 supplemented with 1% BSA were treated with 1:1,000 anti-Fas (human, activating, clone CH11) and 40 μM trova. Different concentrations of MBF, EGTA, ML218, NNC 55-0396 or penfluridol were also added. Anti-Fas, Trova, MBF, EGTA and penfluridol were purchased from Sigma-Aldrich. ML218 and NNC 55-0396 were purchased from Tocris Bioscience. Cell samples were incubated at 37°C in a humidified atmosphere with 5% CO2 for 3 h prior to A5 and TO-PRO-3 staining for flow cytometry analysis.

In *in vitro* infection experiments, 10 μM MBF, 10 μM ML218, 4 μM NNC 55-0396 or 4 μM penfluridol was added into Vero E6 cells following SARS-CoV-2 infection (MOI 0.5) and 1 μg/mL TPCK-treated trypsin. Cells were incubated for 48 h and cells and conditioned media were harvested for ApoBD analysis using flow cytometry using a BD FACSAria™ Fusion flow cytometer.

### ApoBD purification

ApoBDs were purified from conditioned media of ABT-737-treated or SARS-CoV-2-infected Vero E6 cells (thus denoted ABT-ApoBDs and SARS-ApoBDs, respectively) using differential centrifugation as previously described^37^. In brief, apoptotic cell samples were centrifuged at 500 g for 10 min to remove cells. The resultant supernatant was then centrifuged at 3,000 g for 20 min to pellet ApoBDs from smaller EVs. To control for residual extra-ApoBD virus in the SN and/or sticky viruses adhered to SARS-ApoBD, a separate ABT-ApoBD prep was incubated with ApoBD-depleted SARS-CoV-2 supernatant, hence referred to as SN-ApoBDs. All ApoBD samples were washed three times with 1× PBS (Thermo Fisher) and subjected to flow cytometry analysis using ApoBD markers for quality control prior to further assays.

### Caspase-Glo® caspase 3/7 assay

Vero E6 or Calu-3 cells were seeded in 96-well plates at concentration of 1×10^5^ cells/mL for 24 h, followed by infection with 1:2 titration of SARS-CoV-2 MOIs (0.5-0.03125). ABT-737 (1 μM) was included as an apoptosis control. After 48 h, Caspase-Glo^®^ 3/7 reagent (Promega) was added to each well as per manufacturer’s instruction. Cells were incubated for 3 h at 37 °C in a humidified atmosphere with 5% CO2 prior to luminescence reading using a spectrophotometer.

### Microscopy

#### Transmission electron microscopy

Purified ApoBDs were fixed in 3% glutaraldehyde/PBS for 2 hours at room temperature, then embedded in 1.5% low melting point agarose. After washing with 0.1 M cacodylate buffer, the samples were treated with 1% OsO4 (in 0.1 M cacodylate buffer) for 40 minutes at room temperature. Following a buffer wash and an 80% acetone wash, samples were incubated overnight at 4°C in 2% uranyl acetate/80% acetone. On the next day, sample dehydration and resin infiltration was performed as followed: 2 × 10 min with 80% acetone, 2 × 10 min with 90% acetone, 3 × 20 min with 100% acetone, 1 × 90 min with 50% Epon/50% acetone, 1 × 90 min with 75% Epon/25% acetone, and 1 × 90 min with 100% Epon. Fresh 100% Epon with benzyldimethylamine was then added and samples were embedded at 60°C for 72 h. Resin blocks were sliced into 50 nm thick sections using DiATOME 45° Ultra Diamond Knife on a Leica EM UC7 ultramicrotome. These sections were mounted on EM-copper grids with formvar/carbon coating, post-stained in 4% aqueous uranyl acetate and Reynolds’ lead citrate before imaging with a FEI Talos L120C transmission electron microscope.

#### Bright-field imaging

Vero E6 or Calu-3 cells were seeded overnight in 96-well plates at 1×10^5^ cells/mL. On the next day, freshly purified ApoBDs were added into each well (cell:ApoBD ratio of 1:3). After 48 h, cytopathic effect was visualised under the bright-field of a ZOE™ Cell Imager (Bio-Rad).

#### Fluorescence microscopy

At 48 h post-SARS-CoV-2 infection or ApoBD co-incubation, MDMs were washed three times with 1× PBS, followed by fixation (with 2% paraformaldehyde), permeabilisation (with 0.5% saponin) and stained with Hoechst 34580 stain (1 μg/μL; ThermoFisher), Alexa Fluor™ 647 Phalloidin (165 nM; ThermoFisher) and Alexa Fluor® 488-conjugated anti-SARS-CoV-2 Spike S1 subunit antibody (1:200, RnD Systems). Fluorescence imaging was performed using a Zeiss LSM800 laser scanning confocal microscope.

#### Differential interference contrast microscopy

Cells were seeded in an 8-well chambered NuncTM Lab-TekTM II cover glass (Nunc) prior to anti-Fas (1:1,000) and drug treatment. The chambers were pre-treated with 0.01% aqueous poly-L-lysine (Sigma-Aldrich) to enhance cell adherence. Time-lapse differential interference contrast imaging was performed at 37°C with 5% CO2 using spinning disc confocal microscope (Zeiss) with ×63 oil immersion objective. Image processing and analysis was performed using Zeiss imaging software.

#### Calcium imaging

Cells were stained with 2.5 μM cell-permeant Fluo-4 AM (Thermo Fisher)/0.02% pluronic acid F0127 in indicator free media supplemented with 1% BSA at 37°C for 30 min. Cells were then seeded in a poly-L-lysine-coat NuncTM Lab-TekTM II glass chamber followed by apoptosis induction and drug treatment. Calcium influxes post-apoptosis induction was detected at 30 s interval using spinning disc confocal microscope.

### Immunoblotting

Cells or purified ApoBDs were lysed at 4°C using ice-cold 1× cell lysis buffer (BD Biosciences). Protein concentration of the ApoBD lysates was determined using Pierce™ bicinchoninic acid assay (Thermo Fisher). Subsequently, 30 μg of total protein per sample was subjected to reducing and denaturing SDS-PAGE prior to Western transfer onto nitrocellulose membrane (Bio-Rad). After blocking with 5% skim milk in PBS containing 0.1% (v/v) Tween®-20 (PBST), the membrane was probed with antibodies as followed: anti-human PANX1 (1:1,000), anti-human CaV3.2 (1:500), anti-human β-actin (1:1,000) (all rabbit antibodies from Abcam) and horseradish peroxidase-conjugated donkey-anti-rabbit IgG (1:10,000; Cell Signaling Technology) in PBST containing 1% (w/v) BSA. Chemiluminescence detection was achieved using ECL Primer reagents (GE Healthcare, UK).

### RT-qPCR

RNA was isolated from TRIzol samples using Phasemaker tubes (Life Technologies) with chloroform:isoamyl alcohol (49:1 v/v, Sigma-Aldrich) and precipitated with isopropanol (Sigma-Aldrich), 0.3 M sodium acetate (Sigma-Aldrich) and 10 mg/mL linear polyacrylamide (Life Technologies) at -20°C overnight. The precipitated RNA was washed twice with 80% ethanol and then resuspended in distilled H_2_O. RNA sample concentrations were measured using the Qubit 4 fluorometer (ThermoFisher) and the RNA Broad Range assay kit (ThermoFisher). SARS-CoV-2 Nucleocapsid-1 (N1) positive control plasmid (250,000 copies per μL, IDT 10006625) was used to create a standard curve by performing seven two-fold dilutions (up to 1:128). iTaq Universal Probes One-Step Kit (Bio-Rad 1725141) master mix was made up according to manufacturers’ instructions. Master mix was vortexed and 16 μL was added to each well of a 96-well Light cycler plate followed by 4 μL of RNA (standardised to 40 ng per sample), positive control (standards) or negative control (H_2_O). The plate was briefly pulsed in the centrifuge and sealed with a clear PCR plate seal (Roche Life Science). The LightCycler® instrument (Roche) was run using FAM dye as the reporter and the following cycling parameters:

| Step | No. of cycles | Temperature | Time |
| --- | --- | --- | --- |
| Pre-incubation | 1 | 50°C | 10 mins |
|  | 1 | 95°C | 2 mins |
| Two-step amplification | 50 | 95°C | 60 s |
|  |  | 60°C | 30 s |
| Cooling | 1 | 37°C | 30 s |

Data were analysed by performing an absolute quantification against the standard curve using the LightCycler® 96 analysis software (Roche).

*ACE2* and *TMPRSS2* expression were quantified as previously described^52^.

### TCID50 assay

Purified ApoBDs were resuspended in infection media (DMEM containing 1 μg/mL trypsin-TPCK) to 1×10^6^ ApoBDs/mL. Infectious viral titre was measured using TCID50 assay by plating 1:7 serially diluted ApoBD suspension onto confluent layers of Vero E6 cells in replicates of six on 96-well plates. Plates were incubated at 37°C supplied with 5% CO2 for four days before scoring cytopathic effect under light microscope. The TCID50 calculation was performed using the Spearman Karber method^56^.

For murine lung TCID50 assay, left lungs were harvested and stored at −80 °C in serum-free DMEM until further processing. Lung samples were defrosted and homogenised in a Bullet Blender (Next Advance Inc) in 1 mL DME media (ThermoFisher) containing five steel homogenization beads (Next Advance Inc). Lung homogenates were then centrifuged at 10,000 *g* for 5 min at 4°C. Supernatant was retained for TCID50 assay to determine viral load.

### ApoBD and macrophage co-incubation

Human MDMs were prepared by seeding 2.5×10^5^ CD14^+^ monocytes onto 48-well plate in RPMI 1640 media supplemented with 1 mM L-glutamine, 2 mM sodium pyruvate, 10% human adult AB serum and 5 ng/mL GM-CSF. After seven days, MDMs were washed with 1× PBS before co-incubation with freshly-purified ApoBDs (MDM:ApoBD ratio of 1:3, equivalent to SARS-CoV-2 MOI of 3) or infection with SARS-CoV-2 (in the presence of 1 μg/mL trypsin-TPCK, MOI of 3) for 4 h. Infection media or excess ApoBDs were then removed and replaced with fresh RPMI 1640 media supplemented with 5% human adult AB serum, without GM-CSF.

### Cell viability assay

The viability of SARS-CoV-2-infected or ApoBD-treated MDMs after 48 h were determined using tetrazolium-based calorimetric assay as previously described^57^.

### Cytokine bead array

Cytokine released by the SARS-CoV-2-infected or ApoBD-treated MDMs were measured using human cytokine bead array as per manufacturer’s instruction (BD Biosciences). Briefly, conditioned media from were collected at 48 h. A mixture of 2.5 μL each of captured cytokine beads and 15 μL of PE detection reagent was prepared per sample. This bead cocktail (30 μL) was incubated with an equal volume of cytokine standards or conditioned media in a 96-well plate and incubated for 3 hours. After washes by centrifugation (200 g, 5 min, room temperature), samples were resuspended in 80 μL of wash buffer, then analysed using a BD FACSAria™ Fusion flow cytometer. Data analysis was performed using FCAP Array software v.3 (BD Biosciences).

In murine BAL experiments, BAL supernatant after 3,000 *g* centrifugation was retained and subjected to cytokine bead array analysis as described.

#### *In vitro* engulfment assay

The ability of MDMs to efferocytose ApoBDs was quantified using an engulfment assay as previously described^18^. Briefly, macrophages and ApoBDs, pre-stained with CypHer5E NHS Ester (a pH-sensitive engulfment indicator, Sigma-Aldrich) were co-incubated at 1:3 ratio. At 4 hours post-incubation, MDMs were harvested using Accutase® cell detachment solution (STEMCELL Technologies) for flow cytometry analysis a BD FACSAria™ Fusion. For inhibition of efferocytosis, CypHer5E-stained ApoBDs were blocked with BD Pharmingen™ purified recombinant A5 in 1× A5 binding buffer (BD Biosciences; 50 μg for every 1×10^5^ ApoBDs). Alternatively, MDMs were pre-treated with 5 μM cytochalasin D (Thermo Fisher) for 30 min before co-incubation.

### Fluorescence-activated cell sorting

FACS experiments were performed as per engulfment assay but using 6-well plates (CD14^+^ monocytes seeded at 2×10^6^ cells/well). After co-incubation with CypHer5E-stained ApoBDs, MDMs were detached and resuspended in FACS buffer (1× PBS, 1% FBS, 2 mM EGTA). Cell suspensions were strained through 70 μm nylon mesh filters into FACS tubes. ApoBD engulfers and non-engulfers (2×10^5^ each) were separately sorted into complete media-containing collection tubes based on CypHer5E^+^ and CypHer5E^-^ respectively, using a BD FACSAria™ Fusion flow cytometer and FACSDiva 6.1.1 software (BD Bioscience). Sorted MDMs were then pelleted at 500 g for 5 min and plated onto a 48-well plate. After 48 h, MDMs were harvested for flow cytometry and qRT-PCR analyses whilst condition media were collected for human cytokine bead array.

### CRISPR/Cas9 gene editing

Jurkat T cells deficient of CaV3.2 were generated using a CRISPR/Cas9 system as previously described^58^, using gRNA targeting CACNA1H exon 4: 5’-CAGGCCTTTGACGCCTTCAT-3’.

### Mibefradil conditioned media transfer assay

Vehicle control or MBF (10 μM) was added onto Vero E6 cells following SARS-CoV-2 infection (MOI 0.5) and 1 μg/mL TPCK-treated trypsin. After 48 h incubation, conditioned media were collected and centrifuged at 3000 g for 20 min. After decanting supernatant containing small EVs and free viruses, pellet was resuspended in complete RPMI 1640 and transferred on to MDMs for 4 h. Subsequently, MDMs were washed with 1× PBS and further cultured in fresh RPMI 1640 media supplemented with 1 mM L-glutamine, 2 mM sodium pyruvate, 10% human adult AB serum for 48 h.

### BAL flow cytometry analysis

BAL samples were collected from mice upon culling and centrifuged at 3,000 *g* for 20 min to collect cells and ApoBDs for staining with a combination of LiveDead-APC stain (Thermo Fisher), SR-DEVD-FMK and A5-BV605 in staining buffer (1× A5 binding buffer, 2 mM EDTA 2% FCS, 1× PBS). After washing with staining buffer, samples were incubated with Fc block (BD Biosciences) and stained with a surface marker cocktail containing Ly6G-PE-Cy7 (1:300), F4/80-PerCP-Cy5.5 (1:150), CD45.2-AF700 (1:200), CD11C-APC-Cy7 (1:200), Ly6C-BV421 (1:300) and CD11b-V500 (1:300). All antibodies were purchased from BD Biosciences. Following fixation (with 2% paraformaldehyde) and permeabilisation (using 0.5% saponin), samples were incubated with FITC-conjugated anti-viral spike antibodies (1:250, RnD Systems). At the end, SPHERO™ AccuCount beads (Spherotech) were added prior to flow cytometry analysis using a BD LSRFortessa™ Cell Analyser.

### Lung histological analysis

Right lungs were harvested and fixed in 4% paraformaldehyde for 24 h before transferring to 70% ethanol for dehydration. Lung samples were then paraffin embedded and sectioned for routine histology conducted by the WEHI Histology Facility. Slides were scanned using a 3D Histech Brightfiled ×20 with CaseCenter software at 20× magnification.

### Statistical analysis

Data represent mean ± SEM. Statistical significance was determined using one-way ANOVA with a Tukey post-hoc test or Student’s t-test where appropriate. A *p*-value of <0.05 was considered statistically significant. Sample size and number of replicates are specified within the figure legends.

## DATA AVAILABILITY ACKNOWLEDGEMENTS

We thank La Trobe University Bioimaging Platform and the Water and Eliza Hall Institute Flow Cytometry and PC3 Facility as well as Histology Department for their technical supports and services. This work was funded by the National Health and Medical Research Council (NHMRC GNT1173662 (I.P), NHMRC 2025759 (T.K.P)), the Jack Brockhoff Foundation and La Trobe University (T.K.P). A.K.C and D.S are supported by the NHMRC (GNT2020750) and WEHI philanthropy.

## AUTHOR CONTRIBUTIONS

The study was conceived, designed and supervised by T.K.P., A.K.C and I.K.H.P. T.K.P, D.S., B.S., M.D., L.M, D.C.O., G.A.S., O.A., R.T., G.A. performed experiments. K.C.D. and M. Doerflinger contributed to experimental design and discussions. All authors analysed and interpreted the data. The manuscript was written by T.K.P., A.K.C and I.K.H.P., reviewed and approved by all co-authors.

## ADDITIONAL INFORMATION

## ADDITIONAL INFORMATION

## REFERENCES

1. Jackson CB, Farzan M, Chen B, Choe H (2022) Mechanisms of SARS-CoV-2 entry into cells. Nat. Rev. Mol. Cell Biol. 23: 3–20

2. Darif D, Hammi I, Kihel A, El Idrissi Saik I, et al. (2021) The pro-inflammatory cytokines in COVID-19 pathogenesis: What goes wrong? Microb. Pathog. 153: 104799

3. Neufeldt CJ, Cerikan B, Cortese M, Frankish J, et al. (2022) SARS-CoV-2 infection induces a proinflammatory cytokine response through cGAS-STING and NF-κB. Commun. Biol. 5: 45

4. Merad M, Martin JC (2020) Pathological inflammation in patients with COVID-19: a key role for monocytes and macrophages. Nat. Rev. Immunol. 20: 355–362

5. Huang Q, Wu X, Zheng X, Luo S, et al. (2020) Targeting inflammation and cytokine storm in COVID-19. Pharmacol. Res. 159: 105051

6. Hamidzadeh K, Christensen SM, Dalby E, Chandrasekaran P, et al. (2017) Macrophages and the Recovery from Acute and Chronic Inflammation. Annu. Rev. Physiol. 79: 567–592

7. Bost P, Giladi A, Liu Y, Bendjelal Y, et al. (2020) Host-Viral Infection Maps Reveal Signatures of Severe COVID-19 Patients. Cell 181: 1475–1488.e12

8. Chua RL, Lukassen S, Trump S, Hennig BP, et al. (2020) COVID-19 severity correlates with airway epithelium–immune cell interactions identified by single-cell analysis. Nat. Biotechnol. 38: 970–979

9. Zheng J, Wang Y, Li K, Meyerholz DK, et al. (2021) Severe Acute Respiratory Syndrome Coronavirus 2–Induced Immune Activation and Death of Monocyte-Derived Human Macrophages and Dendritic Cells. J. Infect. Dis. 223: 785–795

10. González Pessolani T, Muñóz Fernández de Legaria M, Elices Apellániz M, Salinas Moreno S, et al. (2021) Multi-organ pathological findings associated with COVID-19 in postmortem needle core biopsies in four patients and a review of the current literature. Rev. Española Patol. 54: 275–280

11. Zhang Z, Penn R, Barclay WS, Giotis ES (2022) Naïve Human Macrophages Are Refractory to SARS-CoV-2 Infection and Exhibit a Modest Inflammatory Response Early in Infection. Viruses 14: 441

12. Niles MA, Gogesch P, Kronhart S, Ortega Iannazzo S, et al. (2021) Macrophages and Dendritic Cells Are Not the Major Source of Pro-Inflammatory Cytokines Upon SARS-CoV-2 Infection. Front. Immunol. 12:

13. Thorne LG, Reuschl A, Zuliani‐Alvarez L, Whelan MVX, et al. (2021) SARS‐CoV‐2 sensing by RIG‐ I and MDA5 links epithelial infection to macrophage inflammation. EMBO J. 40:

14. García-Nicolás O, V’kovski P, Zettl F, Zimmer G, et al. (2021) No Evidence for Human Monocyte-Derived Macrophage Infection and Antibody-Mediated Enhancement of SARS-CoV-2 Infection. Front. Cell. Infect. Microbiol. 11:

15. Sefik E, Qu R, Junqueira C, Kaffe E, et al. (2022) Inflammasome activation in infected macrophages drives COVID-19 pathology. Nature 606: 585–593

16. Poon IKH, Lucas CD, Rossi AG, Ravichandran KS (2014) Apoptotic cell clearance: Basic biology and therapeutic potential. Nat. Rev. Immunol.

17. Atkin-Smith GKGK, Miles MAMA, Tixeira R, Lay FTFT, et al. (2019) Plexin B2 Is a Regulator of Monocyte Apoptotic Cell Disassembly. Cell Rep. 29: 1821–1831

18. Tixeira R, Phan TK, Caruso S, Shi BB, et al. (2020) ROCK1 but not LIMK1 or PAK2 is a key regulator of apoptotic membrane blebbing and cell disassembly. Cell Death Differ. 27: 102–116

19. Brock CK, Wallin ST, Ruiz OE, Samms KM, et al. (2019) Stem cell proliferation is induced by apoptotic bodies from dying cells during epithelial tissue maintenance. Nat. Commun.

20. Coleman ML, Sahai EA, Yeo M, Bosch M, et al. (2001) Membrane blebbing during apoptosis results from caspase-mediated activation of ROCK I. Nat. Cell Biol.

21. Sebbagh M, Renvoizé C, Hamelin J, Riché N, et al. (2001) Caspase-3-mediated cleavage of ROCK I induces MLC phosphorylation and apoptotic membrane blebbing. Nat. Cell Biol.

22. Poon IK, Chiu YH, Armstrong AJ, Kinchen JM, et al. (2014) Unexpected link between an antibiotic, pannexin channels and apoptosis. Nature 507: 329–334

23. Moss DK, Betin VM, Malesinski SD, Lane JD (2006) A novel role for microtubules in apoptotic chromatin dynamics and cellular fragmentation. J Cell Sci 119: 2362–2374

24. Schaible UE, Winau F, Sieling PA, Fischer K, et al. (2003) Apoptosis facilitates antigen presentation to T lymphocytes through MHC-I and CD1 in tuberculosis. Nat. Med.

25. Phan TK, Ozkocak DC, Poon IKH (2020) Unleashing the therapeutic potential of apoptotic bodies. Biochem. Soc. Trans. 48: 2079–2088

26. Atkin-Smith GKGK, Duan M, Zanker DJDJ, Loh L, et al. (2020) Monocyte apoptotic bodies are vehicles for influenza A virus propagation. Commun. Biol. 3: 1–14

27. Ganesan M, Natarajan SK, Zhang J, Mott JL, et al. (2016) Role of apoptotic hepatocytes in HCV dissemination: Regulation by acetaldehyde. Am. J. Physiol. - Gastrointest. Liver Physiol.

28. Chu H, Shuai H, Hou Y, Zhang X, et al. (2021) Targeting highly pathogenic coronavirus-induced apoptosis reduces viral pathogenesis and disease severity. Sci. Adv. 7:

29. Ren Y, Shu T, Wu D, Mu J, et al. (2020) The ORF3a protein of SARS-CoV-2 induces apoptosis in cells. Cellular and Molecular Immunology

30. Bader SM, Cooney JP, Pellegrini M, Doerflinger M (2022) Programmed cell death: the pathways to severe COVID-19? Biochem. J. 479: 609–628

31. Morais da Silva M, Lira de Lucena AS, Paiva Júnior S de SL, Florêncio De Carvalho VM, et al. (2022) Cell death mechanisms involved in cell injury caused by SARS‐CoV‐2. Rev. Med. Virol. 32:

32. Liu Y, Garron TM, Chang Q, Su Z, et al. (2021) Cell-Type Apoptosis in Lung during SARS-CoV-2 Infection. Pathogens 10: 509

33. André S, Picard M, Cezar R, Roux-Dalvai F, et al. (2022) T cell apoptosis characterizes severe Covid-19 disease. Cell Death Differ. 29: 1486–1499

34. Varga Z, Flammer AJ, Steiger P, Haberecker M, et al. (2020) Endothelial cell infection and endotheliitis in COVID-19. Lancet

35. Hou Y, Li C, Yoon C, Leung OW, et al. (2022) Enhanced replication of SARS-CoV-2 Omicron BA.2 in human forebrain and midbrain organoids. Signal Transduct. Target. Ther. 7: 381

36. Jiang L, Paone S, Caruso S, Atkin-Smith GKGK, et al. (2017) Determining the contents and cell origins of apoptotic bodies by flow cytometry. Sci. Rep. 7:

37. Phan TK, Poon IKH, Atkin-Smith GK (2018) Detection and isolation of apoptotic bodies to high purity. J. Vis. Exp. 2018:

38. Poon IKH, Parkes MAF, Jiang L, Atkin-Smith GKGK, et al. (2019) Moving beyond size and phosphatidylserine exposure: evidence for a diversity of apoptotic cell-derived extracellular vesicles in vitro. J. Extracell. Vesicles 8:

39. Bader SM, Cooney JP, Sheerin D, Taiaroa G, et al. (2023) SARS-CoV-2 mouse adaptation selects virulence mutations that cause TNF-driven age-dependent severe disease with human correlates. Proc. Natl. Acad. Sci. 120:

40. Atkin-Smith GK, Tixeira R, Paone S, Mathivanan S, et al. (2015) A novel mechanism of generating extracellular vesicles during apoptosis via a beads-on-a-string membrane structure. Nat Commun 6: 7439

41. Phan TK, Fonseka P, Tixeira R, Pathan M, et al. (2021) Pannexin-1 channel regulates nuclear content packaging into apoptotic bodies and their size. Proteomics

42. Levine TB, Bernink PJLM, Caspi A, Elkayam U, et al. (2000) Effect of Mibefradil, a T-Type Calcium Channel Blocker, on Morbidity and Mortality in Moderate to Severe Congestive Heart Failure. Circulation 101: 758–764

43. Labzin LI, Chew KY, Eschke K, Wang X, et al. (2023) Macrophage ACE2 is necessary for SARS-CoV-2 replication and subsequent cytokine responses that restrict continued virion release. Sci. Signal. 16:

44. Junqueira C, Crespo Â, Ranjbar S, de Lacerda LB, et al. (2022) FcγR-mediated SARS-CoV-2 infection of monocytes activates inflammation. Nature 606: 576–584

45. Koutsakos M, Rowntree LC, Hensen L, Chua BY, et al. (2021) Integrated immune dynamics define correlates of COVID-19 severity and antibody responses. Cell Reports Med. 2: 100208

46. Lucas C, Wong P, Klein J, Castro TBR, et al. (2020) Longitudinal analyses reveal immunological misfiring in severe COVID-19. Nature 584: 463–469

47. Phan TKTKTK, Ozkocak DCDC, Poon IKHIKH (2020) Unleashing the therapeutic potential of apoptotic bodies. Biochem. Soc. Trans. 48: 2079–2088

48. Akers JC, Gonda D, Kim R, Carter BS, et al. (2013) Biogenesis of extracellular vesicles (EV): Exosomes, microvesicles, retrovirus-like vesicles, and apoptotic bodies. J. Neurooncol. 113: 1–11

49. Krishnamachary B, Cook C, Kumar A, Spikes L, et al. (2021) Extracellular vesicle‐mediated endothelial apoptosis and EV‐associated proteins correlate with COVID‐19 disease severity. J. Extracell. Vesicles 10:

50. Chu H, Hou Y, Yang D, Wen L, et al. (2022) Coronaviruses exploit a host cysteine-aspartic protease for replication. Nature

51. Salina AC, Dos-Santos D, Rodrigues TS, Fortes-Rocha M, et al. (2022) Efferocytosis of SARS-CoV-2-infected dying cells impairs macrophage anti-inflammatory functions and clearance of apoptotic cells. Elife 11:

52. Sheerin D, Abhimanyu, Peton N, Vo W, et al. (2022) Immunopathogenic overlap between COVID-19 and tuberculosis identified from transcriptomic meta-analysis and human macrophage infection. iScience 25: 104464

53. Phan TK, Lay FT, Hulett MD (2018) Importance of phosphoinositide binding by human βdefensin 3 for Akt-dependent cytokine induction: Immunol. Cell Biol. 96: 54–67

54. Kroon EE, Correa-Macedo W, Evans R, Seeger A, et al. (2023) Neutrophil extracellular trap formation and gene programs distinguish TST/IGRA sensitization outcomes among Mycobacterium tuberculosis exposed persons living with HIV. PLOS Genet. 19: e1010888

55. Jiang L, Tixeira R, Caruso S, Aktin-Smith GK, et al. (2016) Monitoring the progression of cell death and disassembly of dying cells by flow cytometry. Nat. Protoc. 4: 655–663

56. Hierholzer JC, Killington RA (1996) Virus isolation and quantitation. in Virology Methods Manual 25–46

57. Phan TK, Lay FT, Poon IK, Hinds MG, et al. (2016) Human β-defensin 3 contains an oncolytic motif that binds PI(4,5)P2 to mediate tumour cell permeabilisation. Oncotarget 7: 2054–2069

58. Kueh AJ, Herold MJ (2016) Using CRISPR/Cas9 Technology for Manipulating Cell Death Regulators. 253–264

