## Supplementary Figure 1 for "Voltage-gated T-type calcium channel blockers reduce apoptotic body mediated SARS-CoV-2 cell-to-cell spread and subsequent cytokine storm"

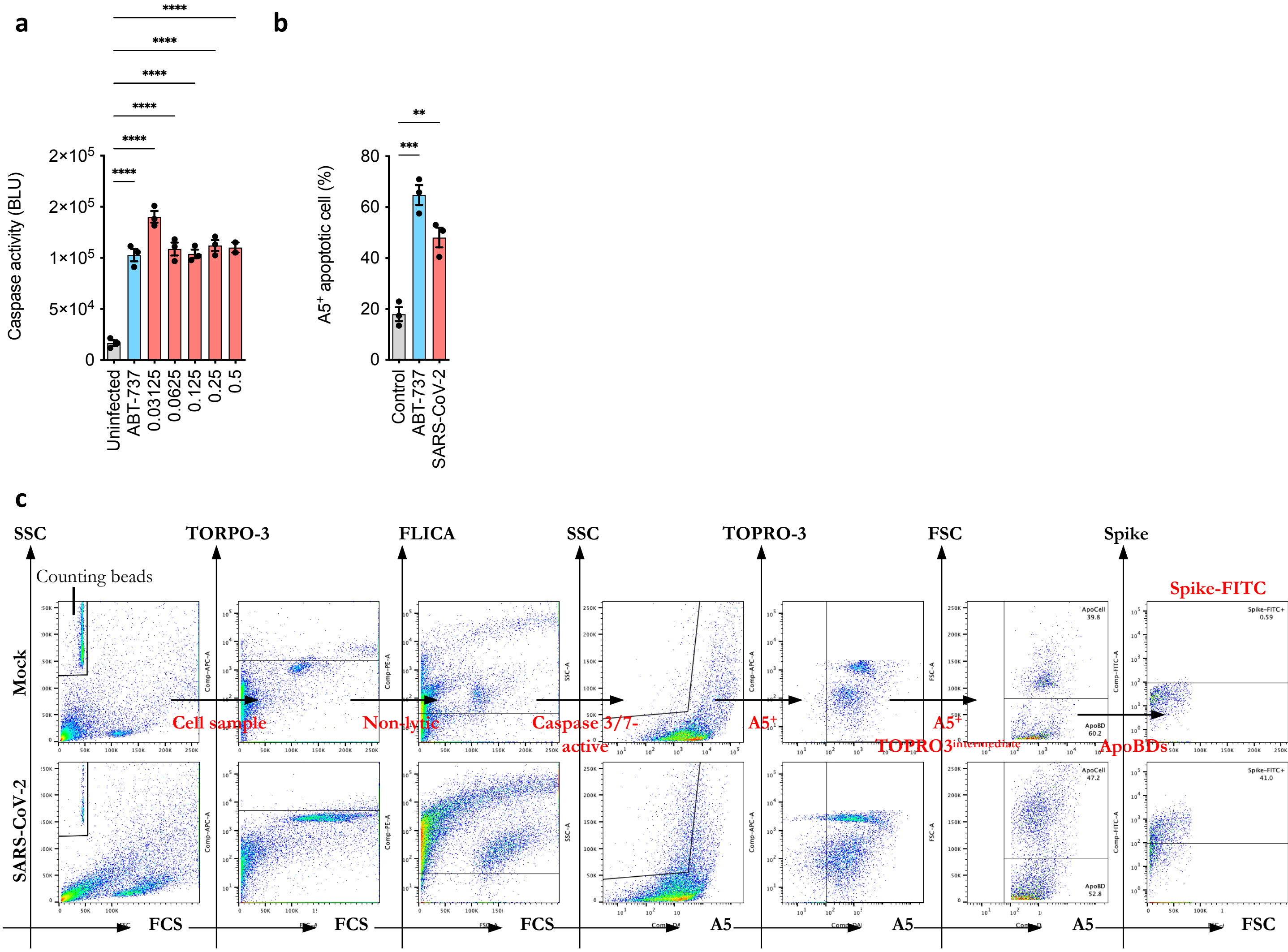


**Supplementary Figure 1. Apoptosis and ApoBD detection upon SARS-CoV-2 infection**

**(a)** Caspase 3/7 activity detected using Caspase 3/7 Glo® bioluminescence assay of SARS-CoV-2-infected Calu-3 lung epithelial cells. **(b)** Detection of apoptotic cells in SARS-CoV-2-infected Vero E6 using flow cytometry analysis with annexin A5 staining. **(c)** Gating strategy for a flow cytometry analysis to detect apoptotic cells and ApoBDs in murine BAL samples, using TOPRO-3 dye, FLICA stain, A5 and viral spike antibody.
