## Supplementary Figure 2 for "Voltage-gated T-type calcium channel blockers reduce apoptotic body mediated SARS-CoV-2 cell-to-cell spread and subsequent cytokine storm"

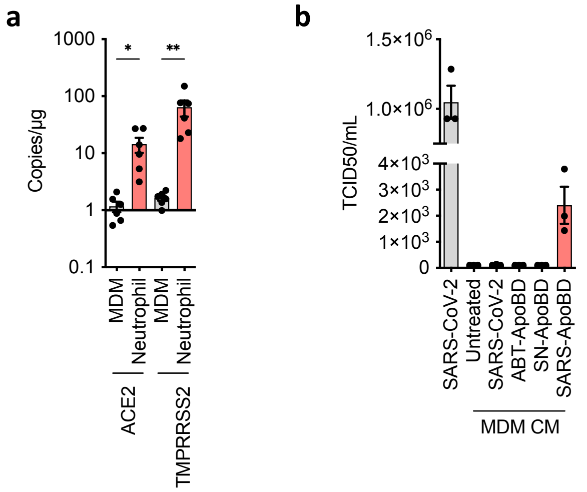


**Supplementary Figure 2. ACE2, TMPRRSS2 expression and SARS-CoV-2 infection in human monocyte-derived macrophages**
**(a)** qPCR analysis to detect ACE2 receptor and TMPRRSS2 protease expression in MDM, compared to SARS-CoV-2-susceptible neutrophils. **(b)** TCID50 assay of conditioned media from MDM culture at 48 h post ApoBD incubation.
