## Supplementary Figure 3 for "Voltage-gated T-type calcium channel blockers reduce apoptotic body mediated SARS-CoV-2 cell-to-cell spread and subsequent cytokine storm"

**a**

| Name | Action | Selectivity | Calcium target | Description |
|---|---|---|---|---|
| (±)-Bay K 8644 | Agonist | L-type calcium channel | | L-type Ca2+ channel agonist |
| Calmidazolium chloride | Inhibitor | Ca2+ATPase | | Potent inhibitor of calmodulin activation of phosphodiesterase; strongly inhibits calmodulin-dependent Ca2+-ATPase |
| NNC 55-0396 | Inhibitor | T-type Ca2+ channels | | Selectiv+K93e T-type calcium channel inhibitor. |
| Mibefradil dihydrochloride | Blocker | T-type Ca2+ channels | | T-type Ca2+ channel blocker |
| SKF 96365 | Inhibitor | TRPC channels; also inhibits store-operated Ca2+ entry | | Selective inhibitor of receptor-mediated and voltage-gated Ca2+ entry |
| Haloperidol | Antagonist | D2/D1 | T-type Ca2+ channels | Dopamine receptor antagonist; antipsychotic |
| Nelfinavir mesylate hydrate | Inhibitor | HIV protease | Inhibits intra-mitochondrial Ca2+ influx | Nelfinavir is a HIV protease inhibitor; antiretroviral; and anti-tumor agent |
| SKF-525A hydrochloride | Inhibitor | Microsomal oxidation | Inhibits transmembrane Ca2+ influx | Inhibitor of microsomal drug metabolism |
| Loperamide hydrochloride | Ligand | Opioid receptor | Blockade of multiple calcium channels | Meperidine congener which binds to opioid receptors; Ca2+ channel antagonist |
| Ifenprodil tartrate | Blocker | Polyamine site NMDA | P-type Ca2+ channel | Blocks the polyamine binding site associated with the NMDA glutamate receptor; neuroprotective |
| BMS-193885 | Antagonist | Y1 | Blocks high-threshold calcium channel | BMS-193885 is a potent, selective Y1 antagonist |

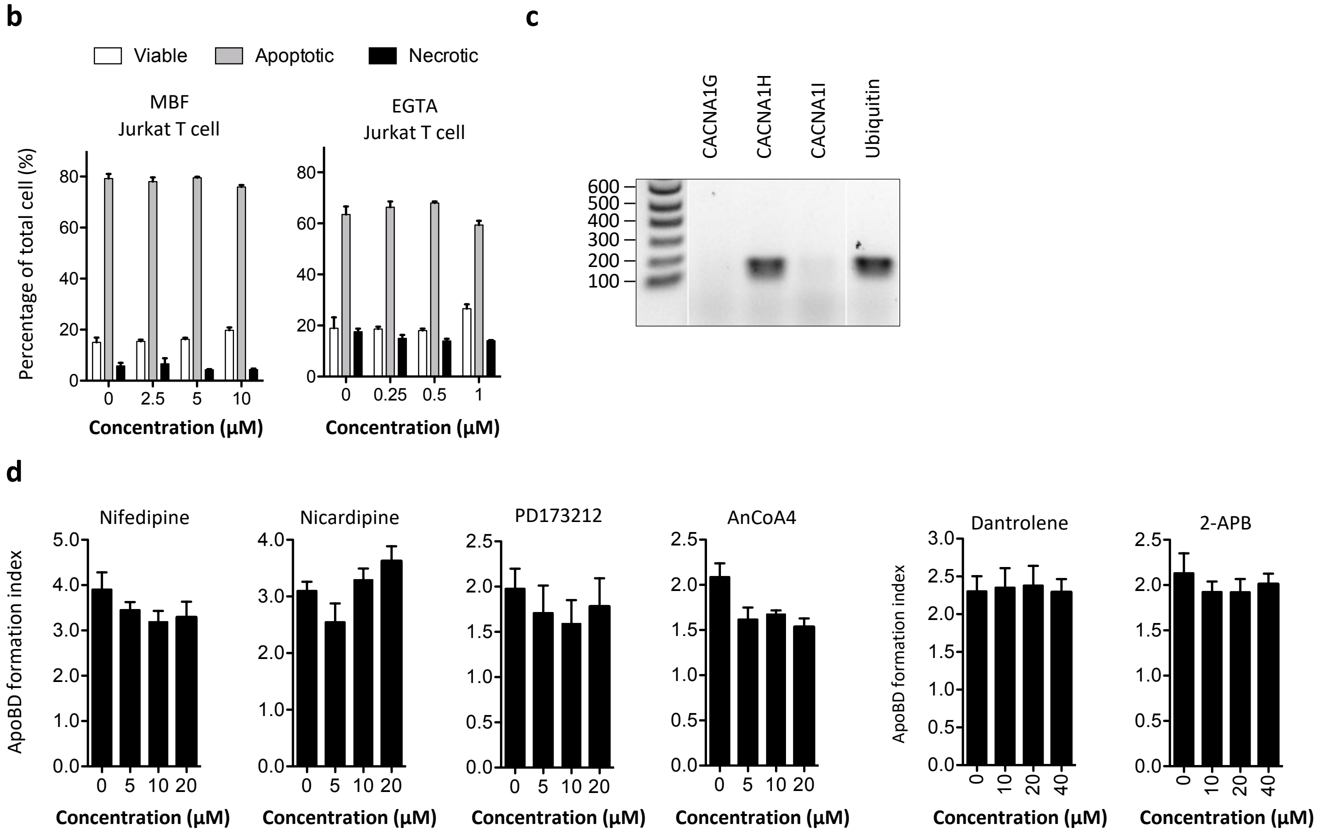


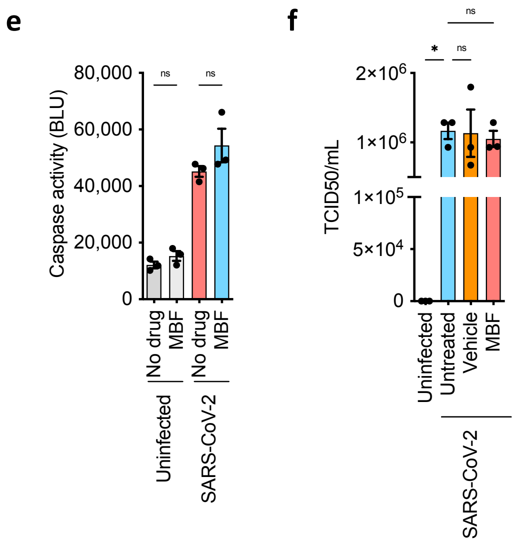


**Supplementary Figure 3. Identification of T-type calcium channel blockers as novel inhibitors of SARS-CoV-2-induced ApoBD formation.**

**(a)** Calcium channel-targeting hits identified through LOPAC® 1280 drug library screening for ApoBD formation inhibitors. **(b)** MBF does not affect level of cell viability, measured by flow cytometry analysis (c) Expression of T-type calcium channels in Jurkat T cells, detected by reverse transcriptase PCR **(d)** ApoBD formation was not impaired by other typical calcium channels including L-type calcium channel (*nifedipine* and *nicardipine*), N-type calcium channel (*PD173212*) or calcium-release-activated calcium channel protein 1 Orai1 (*AnCoA4*), calcium storage release regulated by ryanodine receptor (*dantrolene*) or inositol trisphosphate receptor (*2-aminoethoxydiphenylborane*). **(e)** Level of SARS-CoV-2-induced apoptosis and **(f)** viral infectivity was not affected by MBF treatment in Vero E6 as indicated by Caspase Glo® bioluminescence assay and TCID50 assay.
