## Supplementary Figure 4 for "Voltage-gated T-type calcium channel blockers reduce apoptotic body mediated SARS-CoV-2 cell-to-cell spread and subsequent cytokine storm"

Representative of infected + vehicle

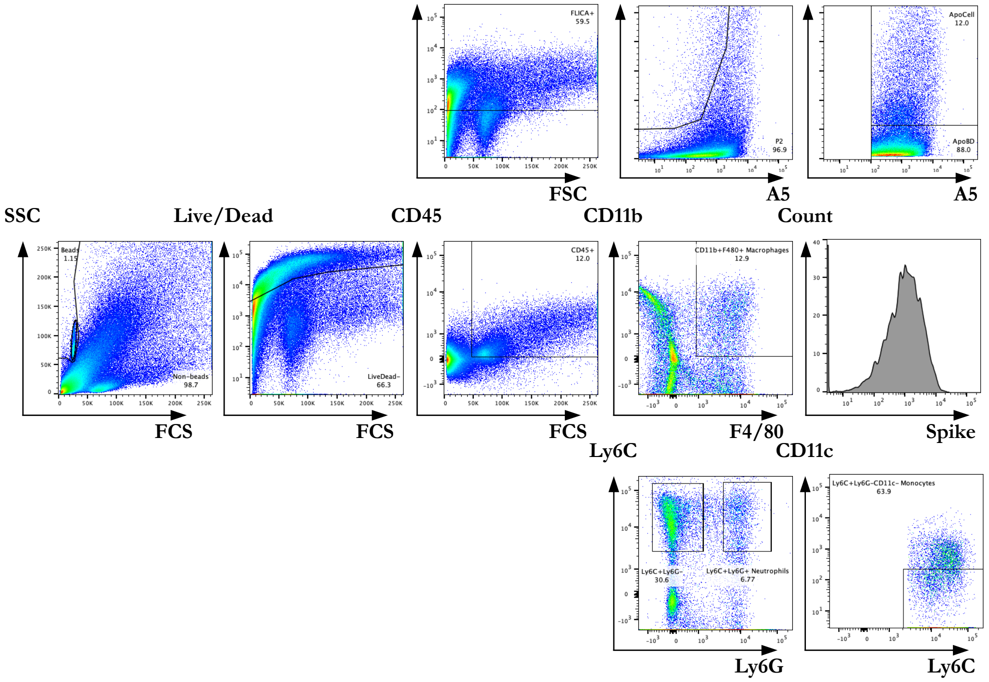


Representative of infected + mibefradil

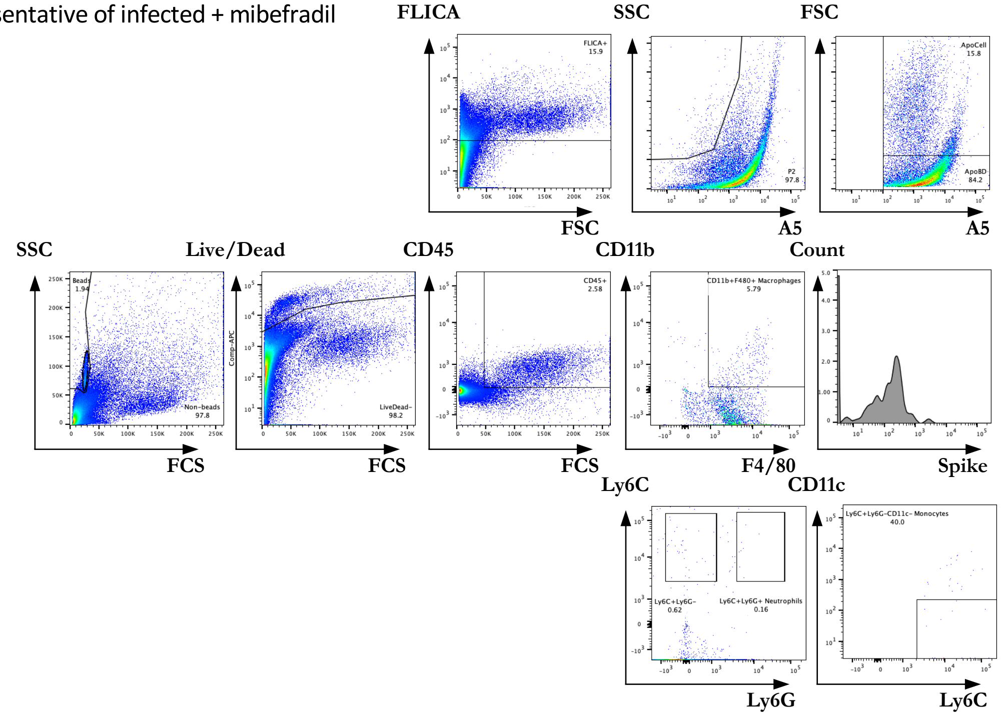


**Supplementary Figure 4. BAL analysis of MBF administration on SARS-CoV-2-infected mice**

Gating strategy for a multicolour flow cytometry analysis to detect ApoBDs, spike⁺ macrophages and immune infiltrates in BAL samples
